# Cyclin D1 regulates the hepatic response to feeding: Evidence for non-cell cycle roles in the liver

**DOI:** 10.64898/2026.05.25.727739

**Authors:** Heng Wu, Jonathan I. Hauser, Na Yang, Nikolai Timchenko, Maggie Klaers, Rahagir Salekeen, Juan C. Manivel, Juan E. Abrahante, Linshan Laux, Matthew J. Yousefzadeh, Michael P. Schonfeld, Irina Tikhanovich, Sayeed Ikramuddin, Satdarshan S. Monga, Oyedele A. Adeyi, Laura J. Niedernhofer, Payel Sen, Matthew S. Gill, Jeffrey H. Albrecht

## Abstract

**Objectives:** Prior studies have shown that cyclin D1 regulates diverse aspects of liver metabolism during cell cycle progression. Interestingly, this protein is induced in hepatocytes by feeding, but its function in modulating hepatic postprandial physiology is poorly characterized. The aim of this study was to evaluate the contribution of cyclin D1 to the hepatic response to feeding and to gain insight into its potential non-proliferative roles in other conditions.

**Methods:** Mice with or without hepatocyte cyclin D1 (D1^fl/fl^ or D1^ΔHep^) were fasted and refed a high-carbohydrate diet. Mouse and human liver in the setting of aging and MASLD were analyzed. The *C. elegans* model was used to evaluate the role of cyclin D1 (CYD-1) in response to overnutrition.

**Results:** Cyclin D1 regulated hepatic gene networks involved in glucose and lipid metabolism, protein synthesis, immune response, and other pathways after feeding. Induction of acute phase response proteins was markedly inhibited in D1^ΔHep^ mice, which was associated with corresponding changes in histone acetylation on key genes. In aged liver, hepatocyte cyclin D1 was induced without associated proliferation; this was markedly pronounced in progeroid *Ercc1*-deficient mice. Cyclin D1 was upregulated in MASLD and diminished with successful treatment. CYD-1 was induced by overnutrition in the intestine of *Caenorhabditis elegans* (which performs metabolic functions similar to liver) and regulates key nutrient-responsive proteins. CYD-1 inhibition prolonged lifespan in this setting.

**Conclusions:** Cyclin D1 regulates nutrient-mediated physiology in the liver and *C. elegans*, indicating that it has unexpected and highly conserved metabolic functions. Further study is warranted to define its role in hepatic disease and aging.

**Highlights:**

- Cyclin D1 is induced in hepatocytes with feeding and broadly regulates hepatic gene expression.
- Acute phase response (APR) and senescence-associated secretory phenotype (SASP) proteins are markedly regulated by cyclin D1.
- Hepatocyte expression of cyclin D1 is substantially upregulated in aging, premature aging, and MASLD without associated proliferation.
- Cyclin D1 (CYD-1) regulates nutrient-mediated signaling and lifespan in response to overnutrition in *C. elegans*.

## 1. INTRODUCTION

The liver plays a central role in metabolism including the response to fasting and feeding [1; 2]. In the fed state, the liver promotes energy storage via glycogen and *de novo* lipid synthesis. In addition, the liver itself undergoes an anabolic response to feeding that includes enhanced protein synthesis and secretion, decreased autophagy, and increased liver size. The complex metabolic adaptations induced by feeding in hepatocytes are controlled by an array of mechanisms including activation and repression of transcription factors, epigenetic modifications, alterations in RNA processing and stability, and translational and post-translational modulation of protein expression. Despite extensive study, the hub proteins that coordinate these diverse regulatory networks are not fully characterized.

Overnutrition and obesity promote increasingly common diseases such as type II diabetes, metabolic syndrome, and metabolic dysfunction-associated steatotic liver disease (MASLD). Furthermore, sustained activation of nutrient-mediated pathways is a driver of aging and cellular senescence, a state characterized by permanent cell cycle arrest, metabolic dysfunction, and the senescence-associated secretory phenotype (SASP) that promotes inflammation and propagates tissue dysfunction [3–5]. Interestingly, the liver is the primary source of many acute phase response (APR) proteins that can also function as SASP, and persistent expression of key APR/SASP proteins can promote metabolic dysfunction and liver diseases [6–10]. Considerable evidence suggests a significant overlap between overnutrition, senescence-related events, APR/SASP expression, aging, inflammation, and metabolic dysfunction in the liver [11–15]. However, key mediators of these overlapping responses remain to be identified.

Cyclin D1 is a cell cycle protein best-known for its role in promoting progression through late G1 phase by binding and activating cyclin-dependent kinase 4 (Cdk4). It is one of the most commonly overexpressed proteins in human cancers [16; 17], acting through Cdk-dependent and -independent mechanisms to induce cell cycle progression and regulate diverse aspects of the malignant phenotype [18–20]. Cyclin D1 plays a pivotal role in physiologic hepatocyte proliferation, such as in the setting of liver regeneration [21–24]. During cell cycle progression, cyclin D1 represses a wide range of homeostatic metabolic pathways in hepatocytes, in part by downregulating the activity of key hepatic transcription factors such as hepatocyte nuclear factor 4α (HNF4α) and peroxisome proliferator-activated receptor α (PPARα) through Cdk4-independent mechanisms [24–26]. For example, in these proliferative models, cyclin D1 diminishes both *de novo* lipogenesis and lipid breakdown (by decreasing lipolysis, lipophagy, autophagy, and fatty acid oxidation) [25–27]; the inhibition of lipid catabolism by cyclin D1 leads to increased steatosis in proliferating hepatocytes [27]. Conversely, cyclin D1 promotes biosynthetic pathways in hepatocytes including glycolysis, the pentose phosphate pathway, and purine and pyrimidine synthesis [28]. Thus, during hepatocyte cell cycle progression, cyclin D1 regulates a broad portfolio of metabolic functions. Paradoxically, prior studies have found that sustained cyclin D1 expression in other cell types can promote cellular senescence through poorly defined mechanisms [3; 29–31], but its potential roles in the liver in aging and disease are not well-studied.

Expression of cyclin D1 in hepatocytes is also induced by refeeding after a period of fasting [24; 32; 33]. Hepatocyte-specific knockout (KO) of cyclin D1 enhances glycogen synthesis [24] and gluconeogenesis [32; 33] in the fed state, indicating that this protein represses key aspects of homeostatic glucose metabolism. Given the wide range of metabolic functions regulated by cyclin D1 during the cell cycle in hepatocytes and other cells [18–20; 24–28; 32–35], we surmised that it may broadly affect postprandial hepatic function independently of cell proliferation. Our findings indicate that hepatocyte cyclin D1 expression regulates diverse pathways in response to feeding, including the induction of APR/SASP proteins. Furthermore, cyclin D1 is elevated in hepatocytes in MASLD and in both physiologic and accelerated aging without a proportionate increase in proliferation. Cyclin D1 (CYD-1) is induced, regulates nutrient-mediated signaling, and shortens lifespan in *C. elegans* in the setting of overnutrition, suggesting that these effects are highly conserved in evolution.

## 2. RESULTS

### 2.1 Cyclin D1 broadly regulates hepatic gene expression in response to feeding

To investigate how cyclin D1 regulates hepatocyte function in response to feeding, we used cyclin D1^fl/fl^ mice [36] for experiments to acutely delete this protein from hepatocytes *in vivo* one week prior to experiments using an established vector that specifically targets the Cre recombinase to these cells (AAV8-TBG-Cre [37; 38]). Mice were then fasted overnight and then refed for 24 h with a high carbohydrate diet (HCD) as described in prior studies [24; 26]. As previously noted [24], western blot demonstrates that cyclin D1 protein is induced by feeding in mice treated with a control vector (GFP), which is eliminated in mice with acute hepatocyte-specific cyclin D1 KO (D1^ΔHep^) mice (Fig. 1A).

**Figure 1.**
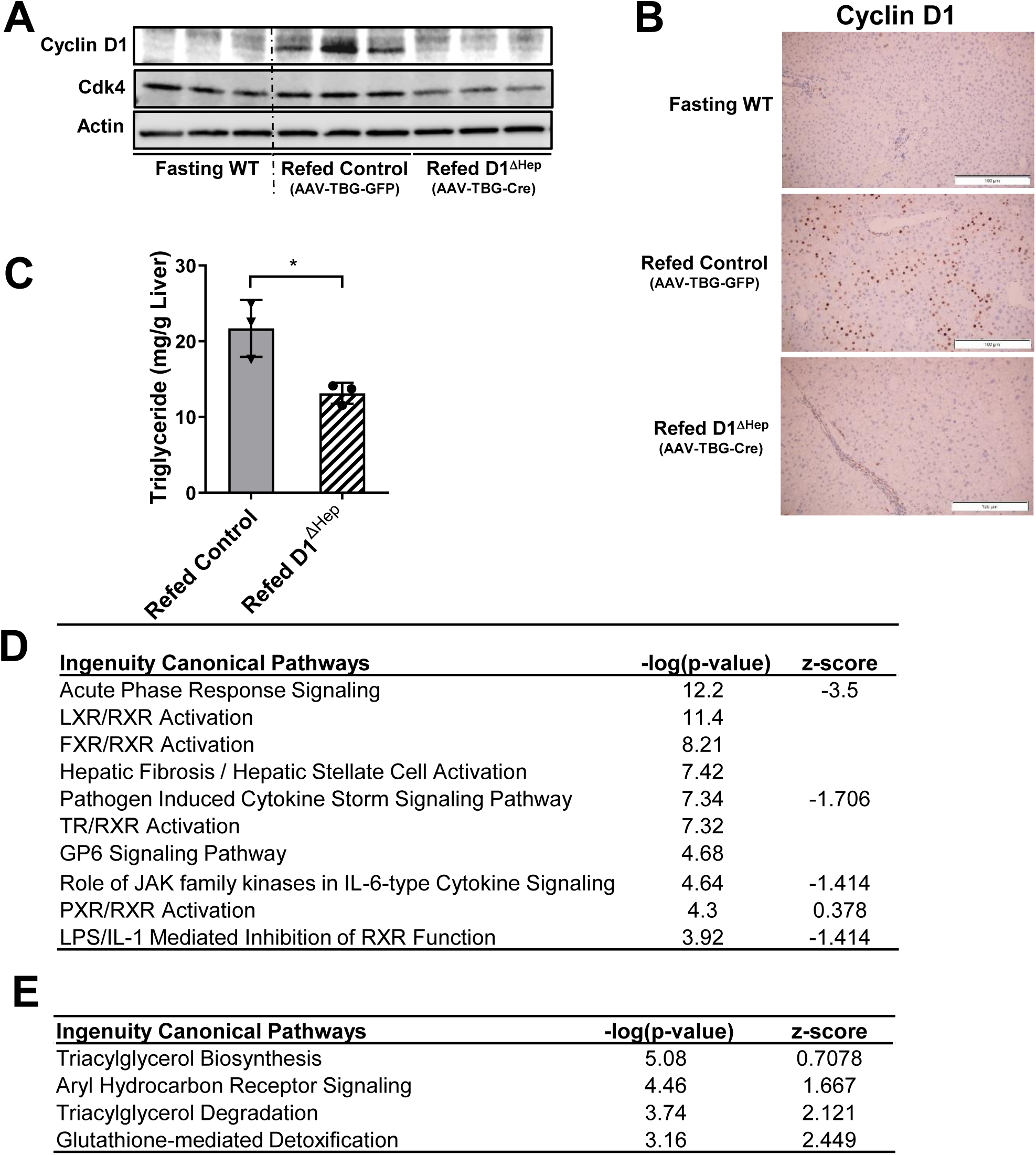
Regulation of hepatic gene expression by cyclin D1 after refeeding. Liver tissue was collected from wild-type (cyclin D1^fl/fl^) after an overnight fast or from control mice (cyclin D1^fl/fl^ with AAV8-TBG- GFP) and D1^ΔHep^ mice (cyclin D1^fl/fl^ with AAV8-TBG-Cre) that were fasted and refed for 24 hours with a high carbohydrate diet. (**A**) Western blot of cyclin D1 and Cdk4 in fasting and refed liver. (**B**) Immunohistochemistry for Cyclin D1. (**C**) Hepatic triglyceride levels. (**D**) The most highly regulated canonical pathways in refed D1^ΔHep^ compared to refed control liver from IPA analysis of RNA-seq data. (**E**) Selected metabolic pathways upregulated in refed D1^ΔHep^ liver.

As shown in Fig. 1B and ref. [24], immunohistochemistry (IHC) demonstrates that refeeding induces cyclin D1 expression in hepatocytes in Zone 2 of the hepatic lobule. The histology also demonstrates enlarged clear cytoplasms in refed mice, which are dramatically larger in the D1^ΔHep^ mice, reflecting a 3-fold increase in hepatocyte glycogen in the absence of cyclin D1 (rather than steatosis) [24]. Refed D1^ΔHep^ mice had lower hepatic triglyceride levels (Fig. 1C), consistent with our prior studies showing that cyclin D1 induces steatosis by decreasing several steps of lipid catabolism [25–27]

We performed RNA-seq on livers isolated from three groups of mice: Fasting wild-type (WT, cyclin D1^fl/fl^), refed control mice (cyclin D1^fl/fl^ with AAV8-TBG-GFP), and refed D1^ΔHep^ mice. Using an expression cut-off of +/- 1.5-fold, 438 transcripts were downregulated and 534 upregulated in refed D1^ΔHep^ livers compared to refed control (Suppl. Table S1). The transcriptomic data were analyzed using Ingenuity Pathway Analysis (IPA), and the most highly affected Canonical Pathways are shown in Fig. 1D and Suppl. Table S2. Unexpectedly, the most highly regulated pathway was Acute Phase Response Signaling, which was markedly downregulated by hepatocyte-specific cyclin D1 knockout (z-score −3.5). Numerous other pathways related to inflammation were also downregulated (Fig. 1D). Conversely, pathways predicted to be enhanced in D1^ΔHep^ liver included a number of metabolic pathways related to homeostatic liver function (Fig. 1E), as predicated by our prior studies in hepatocyte proliferation models [24–27; 34]. These data indicate that cyclin D1 regulates multiple pathways in response to feeding that are not obviously related to cell cycle progression.

We noted that 67% (292 out of 438) of downregulated genes and 53% (284 out of 534) of upregulated genes in refed D1^ΔHep^ liver (relative to control refed mice) were similarly regulated in fasting mice, suggesting that cyclin D1 knockout in fed hepatocytes has effects overlapping with the fasting response. To provide further insight into its effects, we also used our prior transcriptomic data from mice with acute cyclin D1 overexpression in the liver of mice fed *ad lib* [34]. This prior analysis used gene chip technology, which represents substantially fewer transcripts than RNA-seq. Among genes shared on these analyses, there was substantial overlap with transcripts up- or downregulated in refed D1^ΔHep^ livers that were oppositely regulated by cyclin D1 overexpression (Suppl. Table S3).

We have previously shown that overexpression of cyclin D1 in hepatocytes inhibits the transcriptional activity of the carbohydrate response element binding protein (ChREBP, gene name Mlxipl) which is induced in the liver by feeding [26; 39]. Consistent with this, in refed D1^ΔHep^ livers, ChREBP/Mlxipl was predicted to be most upregulated transcription factor in the IPA Upstream Regulator analysis (Suppl. Fig. S1A), and numerous ChREBP-regulated transcripts were significantly more upregulated absence of cyclin D1 (Suppl. Fig. S1B). Similarly, ChREBP (which promotes its own expression [2; 39]) and its key lipogenic target acetyl-Coenzyme A carboxylase α (Acaca) were increased by western blot (Suppl. Fig. S1C). In our prior study, we found that wild-type cyclin D1, but not the cyclin D1-KE point mutant that does not activate Cdk4, inhibited ChREBP transcriptional activity, suggesting that the cyclin D1/Cdk4 kinase regulates ChREBP [26]. We now find that recombinant cyclin D1/Cdk4 phosphorylates ChREBP *in vitro* (Suppl. Fig. S1D), suggesting a possible mechanism of inhibition. Although detailed mechanisms remain to be established, these studies further indicate that physiologic induction of cyclin D1 by refeeding regulates hepatic metabolic function.

### 2.2. Cyclin D1 promotes hepatic expression of APR/SASP and immunoregulatory proteins

In response to acute and chronic systemic stresses, hepatocytes synthesize and secrete high levels of APR proteins (some of which are also SASP) that have diverse local and systemic effects [6–10]. Interestingly, prior studies have also shown that feeding induces hepatic genes and proteins involved in the APR in animal models and humans [40–43]. Accordingly, in the regimen used here, refeeding substantially induced APR/SASP transcripts in control liver, which were markedly diminished in D1^ΔHep^ mice (Fig. 2A).

**Figure 2.**
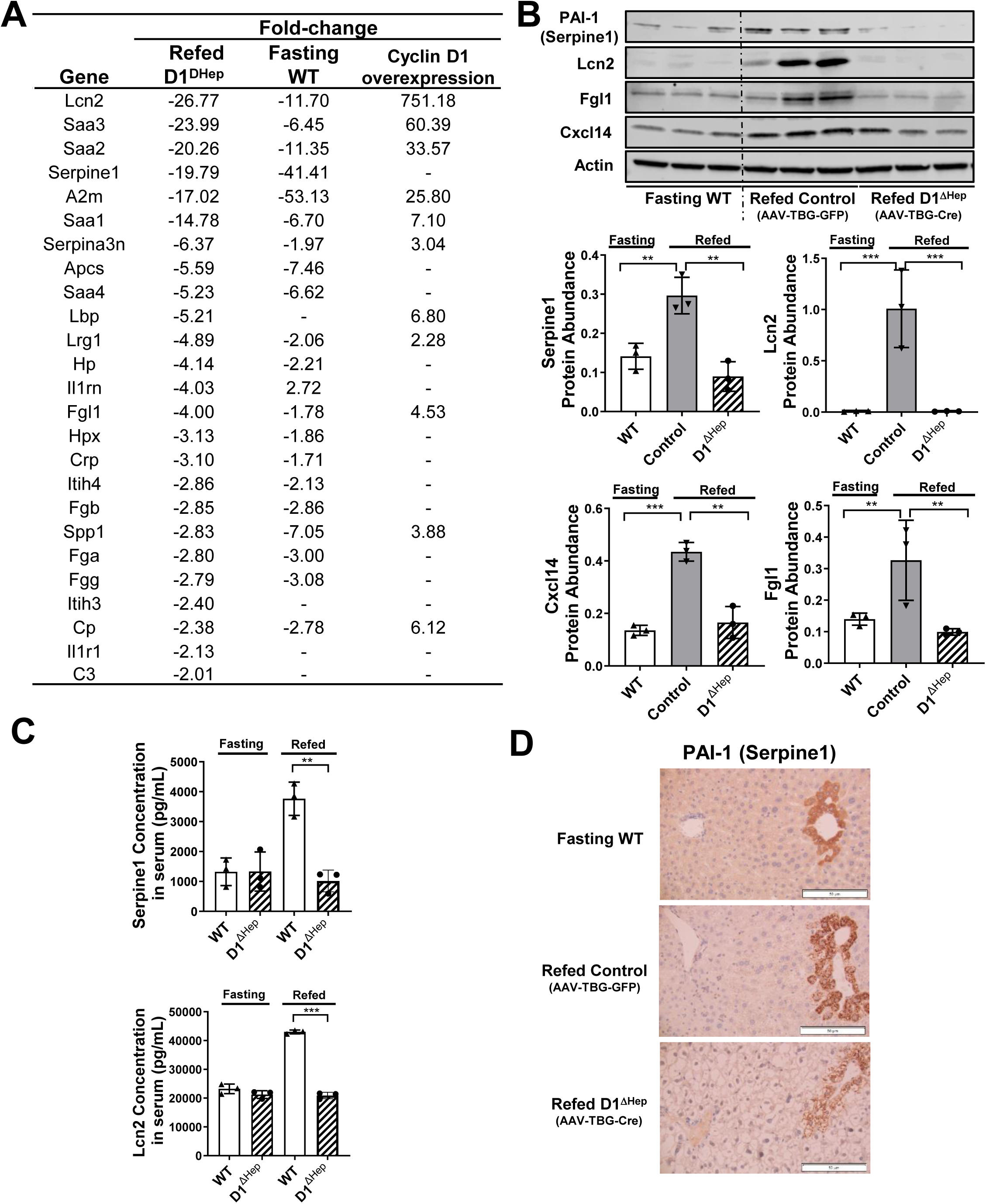
Cyclin D1 regulates hepatic APR/SASP expression. (**A**) Expression of APR transcripts in refed D1^ΔHep^ liver (first column) and fasting liver (second column) compared to refed control liver. The third column shows hepatic expression of transcripts following acute cyclin D1 overexpression (from a prior gene array study [34]). (**B**) Western blot of the indicated proteins. Quantification of protein expression (normalized to actin) is shown. (**C**) Serum levels of Lcn2 and PAI-1 (Serpine1) in the fasting and refed conditions. (**D**) Immunohistochemistry of PAI-1 (Serpine1).

For example, lipocalin 2 (Lcn2) is a multifunctional protein that was markedly induced by feeding and was the most highly downregulated APR/SASP transcript in refed D1^ΔHep^ mice (Fig. 2A). Notably, in a prior study [34], we found that Lcn2 was the most upregulated gene following short-term cyclin D1 expression in the liver (Fig. 2A). The hepatic expression of this protein paralleled the mRNA findings (Fig. 2B). Serum levels of Lcn2 were not affected by hepatocyte cyclin D1 knockout in the fasting state, but its induction with refeeding was ablated in D1^ΔHep^ serum (Fig. 2C). These findings indicate that under the conditions examined here and in our prior study [34], cyclin D1 is both necessary and sufficient for the induction of high-level Lcn2 expression in the liver.

Serpine1, also known as plasminogen activator inhibitor-1 (PAI-1), is an extensively-studied SASP and an APR protein in liver [44–48]. Its expression was induced by refeeding in control but not D1^ΔHep^ liver (Fig. 2A-B). As was the case with Lcn2, the increase in serum PAI-1 after feeding was dependent on hepatocyte expression of cyclin D1 (Fig. 2C). IHC showed that PAI-1 was expressed in the cytoplasm of Zone 3 hepatocytes surrounding the central vein, which was markedly diminished in D1^ΔHep^ liver (Fig. 2D). In fasting mice, PAI-1 expression was relatively uniform in the cytoplasm, whereas there were prominent cytoplasmic PAI-1-positive inclusions with refeeding in control liver, possibly related to the secretory apparatus.

Two other examples of secreted proteins regulated by cyclin D1 were fibrinogen-like protein 1 (Fgl1) and C-X-C Motif Chemokine Ligand 14 (Cxcl14), which were upregulated by feeding and downregulated in D1^ΔHep^ liver (Fig. 2A-B). Fgl1 is primarily synthesized in hepatocytes and has diverse functions, and can play a pathologic role in MASLD, insulin resistance, and immune evasion in cancer [49–52]. Cxcl14 is a cytokine and SASP that can promote liver injury [53]. Cumulatively, the data in Fig. 2 indicate that under these conditions, cyclin D1 substantially promotes the expression of proteins secreted from hepatocytes that have been associated with the APR as well as aging, senescence, and disease.

### 2.3. Cyclin D1 regulates epigenetic modifications in the liver after feeding

Prior studies have shown that cyclin D1 can regulate gene transcription through modification of histone proteins by affecting the recruitment of coregulators such as histone acetylase p300 and histone deacetylase (HDAC) to promoters [54]. In Fig. 3A, cyclin D1 was immunoprecipitated from nuclear extracts and subjected to western blot with p300. The induction of cyclin D1 by refeeding led to its binding to p300, as has been shown in cancer models [54], compatible with the concept that physiologically-expressed cyclin D1 serves a molecular scaffold protein for chromatin modifying enzymes in the liver.

**Figure 3.**
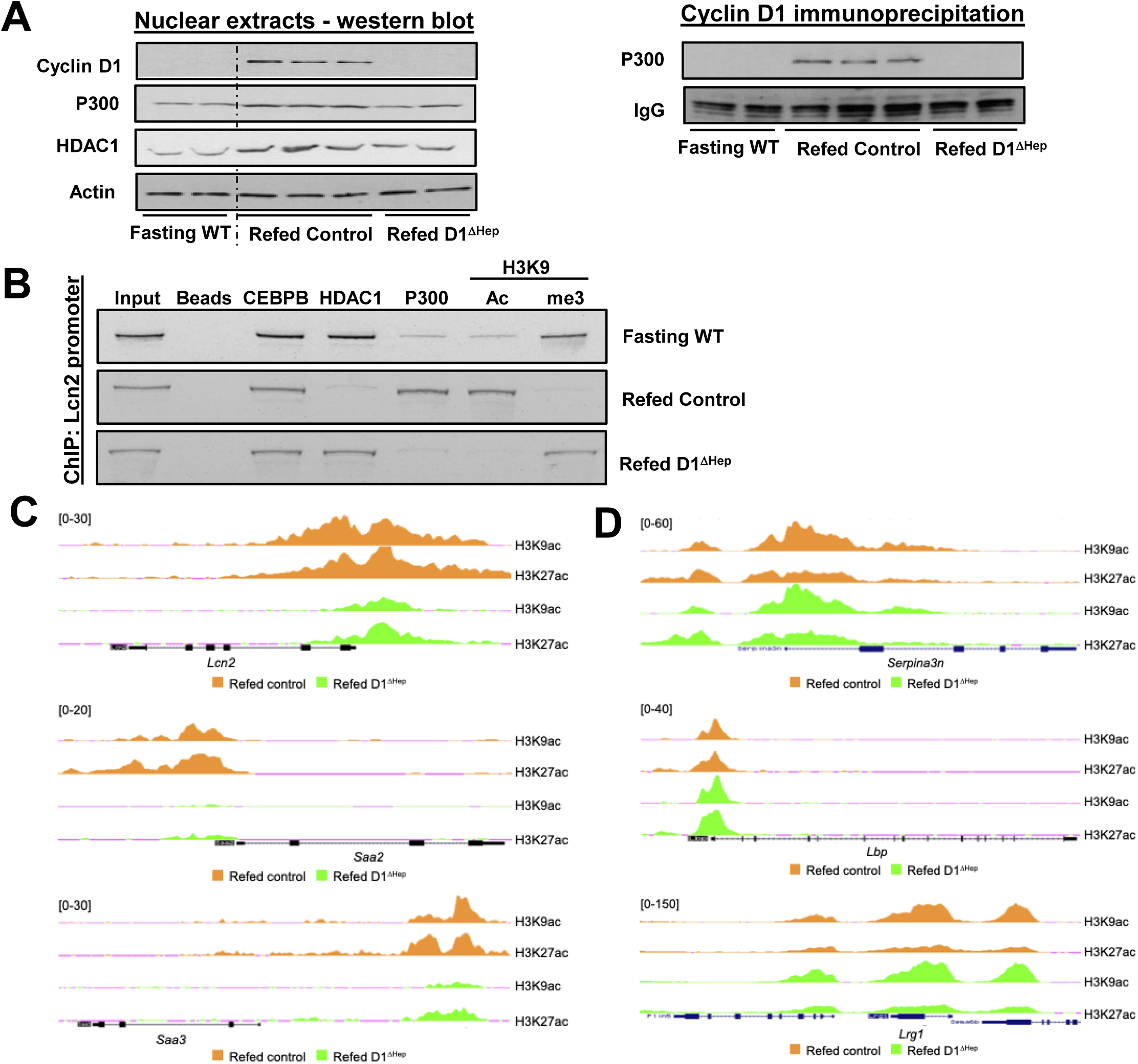
Epigenetic regulation by cyclin D1 in refed liver. (**A**) Western blot of cyclin D1, p300, and HDAC1 in nuclear extracts (left). Western blot of p300 following cyclin D1 immunoprecipitation from nuclear extracts (right). (**B**) Chromatin immunoprecipitation (ChIP) from DNA isolated from liver tissue using the indicated antibodies followed by RT-PCR of a portion of the *Lcn2* promoter. (**C**) ChIP-seq tracings of H3K9ac and H3K27ac occupancy of the genomic regions of *Lcn2*, *Saa2*, and *Saa3*. (**D**) ChIP-seq tracings showing the genomic regions of *Serpina3n*, *Lbp*, and *Lrg1*.

To examine this further, DNA isolated from liver was subjected to chromatin immunoprecipitation (ChIP) with the antibodies shown in Fig. 3B, followed by PCR of a portion of the *Lcn2* promoter. In fasting liver, there was significant binding of HDAC1 and minimal binding of p300 to this promoter. With refeeding in control mice, these patterns were reversed, with diminished HDAC1 and increased p300 binding, along with increased H3K9 acetylation that is generally associated with increased transcription. On the other hand, trimethylation (me3) of H3K9, which is generally associated with transcription repression, was increased in fasting liver and reduced in refed control mice. Importantly, each of these epigenetic changes was absent in refed D1^ΔHep^ liver. Thus, cyclin D1-mediated regulation of histone modifications likely plays a key role in the robust induction of Lcn2 in the liver after feeding. These data also further suggest that cyclin D1 knockout in this setting replicates aspects of the fasting state.

To gain a better understanding of the epigenetic regulation of APR/SASP genes, we performed ChIP-sequencing (ChIP-seq) for H3K9ac and H3K27ac in refed control and D1^ΔHep^ liver. As is shown in Fig. 3C, the three most downregulated APR transcripts in D1^ΔHep^ liver (*Lcn2*, *Saa2*, and *Saa3*) showed a corresponding decrease in histone acetylation, suggesting that cyclin D1 promotes transcription of these genes through epigenetic modifications. However, other APR genes that were inhibited (*Serpina3n*, *Lbp*, and *Lrg1*) showed no significant change in these histone modifications (Fig. 3D), suggesting that other transcriptional or posttranscriptional mechanisms were involved. These data support the concept that cyclin D1 promotes the expression of some, but not all, APR/SASP genes in the liver through epigenetic mechanisms.

Prior studies have suggested that the transcription factor CCAAT/enhancer-binding protein beta (CEBPB) can promote the expression of key APR and SASP proteins [55–57]. Based on the IPA analysis of the RNA-seq data, CEBPB transcriptional activity was predicted to be repressed in refed D1^ΔHep^ liver (Suppl. Fig. S1A). Furthermore, studies in breast cancer cells indicate that cyclin D1 can promote CEBPB activity [58; 59]. To examine the role of this transcription factor in the refeeding model, we used CEBPB^fl/fl^ mice to delete this protein one week prior to experiments using AAV8-TBG-Cre to create hepatocyte-specific knockout (CEBPB^ΔHep^) mice and subjected them to the fasting-refeeding regimen. Surprisingly, based on RNA-seq, there was essentially no difference in the response to refeeding in CEBPB^ΔHep^ liver compared to control (ADV-TBG-GFP treated) mice. *Cebpb* expression was reduced to 0.06-fold compared to control, and *A530013C23Rik* was increased 660-fold, but no other significant changes in gene expression were noted. These results suggest that in contrast to cyclin D1, CEBPB does not have a substantial effect on the hepatic response to feeding in this model. Another possibility is that because CEBPB was deleted from hepatocytes only one week earlier, changes in epigenetic determinants and chromatin accessibility modulated by this protein still persisted, and thus longer-term knockout of this protein may have distinct effects. Additionally, since another member of CEBP family, CEBPA, binds to similar DNA sequences, it is possible that this protein plays a compensatory role in CEBPB^ΔHep^ liver. CEPBP may also play a more prominent function in nonparenchymal cells in the liver (such as immune cells). Further studies are required to unravel the potential role of CEBPB in the hepatocyte response to feeding, but these findings suggest CEBPB is not required to modulate the induction of APR/SASP by cyclin D1.

### 2.4. Hepatocyte cyclin D1 is induced independently of cell cycle progression in aging and MASLD

Our data indicating that cyclin D1 induces expression of SASP proteins led us to question whether it may play a role in liver aging or diseases such as MASLD independently of cell proliferation. Cyclin D1 expression was upregulated in hepatocytes in aged liver (Fig. 4A-B). It was predominantly in Zone 2 and showed a serpiginous pattern throughout the liver, suggesting that contiguous clusters of hepatocytes express this protein (Fig. 4B), and this was not associated with evidence of proliferation (such as ph-Histone H3 expression (Suppl. Fig. S2A)). Aged mouse liver also showed an increase in some of the APR/SASP proteins that are cyclin D1-dependent in refed liver (Fig. 4A). In a small set of histologically normal human livers, cyclin D1 was similarly upregulated in aged liver in hepatocytes (Fig. 4C and Suppl. Fig. S3A). Notably, we did not see evidence of hepatocyte proliferation based on ph-Histone H3 staining (Suppl. Fig. S3A), which is consistent with prior studies showing diminished hepatocyte proliferation in aging [13].

**Figure 4.**
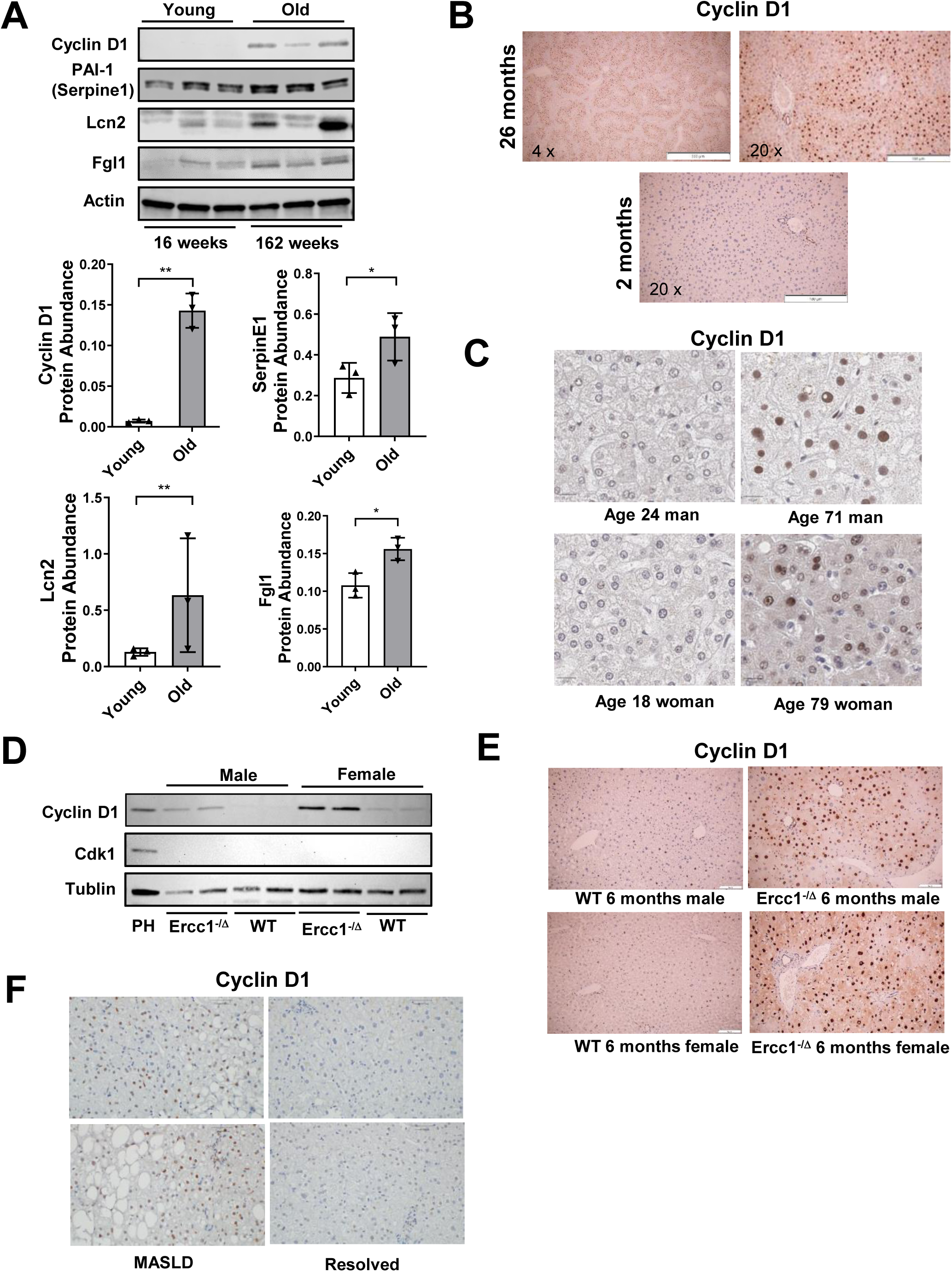
Expression of hepatocyte cyclin D1 in aging and MASLD. (**A**) Western blot of liver from young (4 mo) and old (27 mo) mice. (**B**) IHC for cyclin D1 in old (low and high power) and young mouse liver. (**C**) Cyclin D1 expression in young and old human liver. (**D**) Western blot of Ercc1-deficient (*Ercc1*^-/Δ^) and corresponding WT mouse liver. (**E**) IHC for cyclin D1 in WT and Ercc1-deficient mouse liver. (**F**) IHC for cyclin D1 in liver from two MASLD patients before and after successful treatment with vertical sleeve gastrectomy.

We further examined accelerated aging in Ercc1-deficient (*Ercc*^-/Δ^) mice, which have impaired DNA repair and is a model of the human XFE progeria syndrome [60]. These mice display marked liver dysfunction that plays a role in shortening their lifespan [61]. By western blot, there was substantial induction of cyclin D1 without a corresponding increase in Cdk1, a marker of proliferation in the liver (Fig. 4D). At six months of age, the large majority of Ercc1-deficient hepatocytes expressed cyclin D1 (Fig. 4E) without proliferation as determined by ph-Histone H3 staining (Suppl. Fig. S2A). As previously described [61], Ercc1-deficient liver developed progressive histologic abnormalities including hepatocyte cell death, anisocytosis, dysmorphic cytoplasm, and decreased lobular size. Notably, cyclin D1 expression in *Ercc*^-/Δ^ liver was similar to that seen regenerating liver in young mice 42 h after partial hepatectomy, which show robust hepatocyte proliferation (Fig. 4D and Suppl. Fig. S2A-B) [24]. These studies demonstrate that the marked induction of cyclin D1 in these aging models is uncoupled from cell cycle progression, suggesting that this protein plays a distinct role in this setting.

Prior studies have found enhanced expression of cyclin D1 in MASLD and its more aggressive variant metabolic dysfunction-associated steatohepatitis (MASH) [62–64]. In specimens of human MASLD/MASH, we noted numerous cyclin D1-positive hepatocytes in Zone 2 without corresponding evidence of significant hepatocyte proliferation as determined by ph-Histone H3 or Ki-67 staining (Fig. 4F and Suppl. Fig. S3B). In contrast, in a patient with severe autoimmune hepatitis, which can display both hepatocyte cell death and regeneration [65], there was evidence of both cyclin D1 expression and proliferation (Suppl. Fig. S3B). In paired biopsies, patients with resolved MASLD/MASH after vertical sleeve gastrectomy surgery and weight loss no longer had hepatocyte cyclin D1 expression (Fig. 4F). While the studies in Fig. 4 do not establish a causative role for cyclin D1 in aging or MASLD/MASH pathophysiology, they provide a clinical correlate to support the concept that this protein plays a non-proliferative (and potentially deleterious) role in hepatic metabolism.

### 2.5. Cyclin D1 (CYD-1) plays a role in nutrient-mediated signaling and lifespan in *C. elegans*

The nematode *Caenorhabditis elegans* has been extensively studied as a model of fundamental processes in metabolism, aging, and other aspects of biology. The intestine of *C. elegans* carries out metabolic functions equivalent to the liver and responds similarly to nutritional status [66]. Cyclin D1 has a single ortholog in worms (CYD-1) [67]. To examine whether cyclin D1’s role in feeding-mediated physiology is conserved between species, we created a strain of *C. elegans* linking mScarlet to the endogenous CYD-1 protein, which allows for assessment of its expression in different organs.

Glucose treatment is a model of overnutrition in worms and leads to reduced lifespan [68–70]. CYD-1 expression was increased in the intestine of L4 worms grown on 100mM glucose (Figure 5A) indicating that it is induced by overnutrition similarly to cyclin D1 in carbohydrate-refed mouse liver. Using existing GFP strains [71; 72], we examined the expression of two nutrient-mediated genes that promote lifespan in *C. elegans*. NHR-49 is a well-characterized nuclear receptor transcription factor that regulates the response to starvation [73; 74], and shares homology with mammalian HNF4α and PPARα [75], which we have shown to be inhibited by cyclin D1 in hepatocytes [24–26]. ATGL (ATGL-1) is a pivotal lipase that is also inhibited by cyclin D1 in the liver [27]. As is shown in Fig. 5B-C, CYD-1 knockdown with RNAi increased the intestinal expression of NHR-49::GFP and ATGL-1::GFP in L4 animals. These data indicate cyclin D1/CYD-1 has effects on key metabolic proteins that are highly conserved between species.

**Figure 5.**
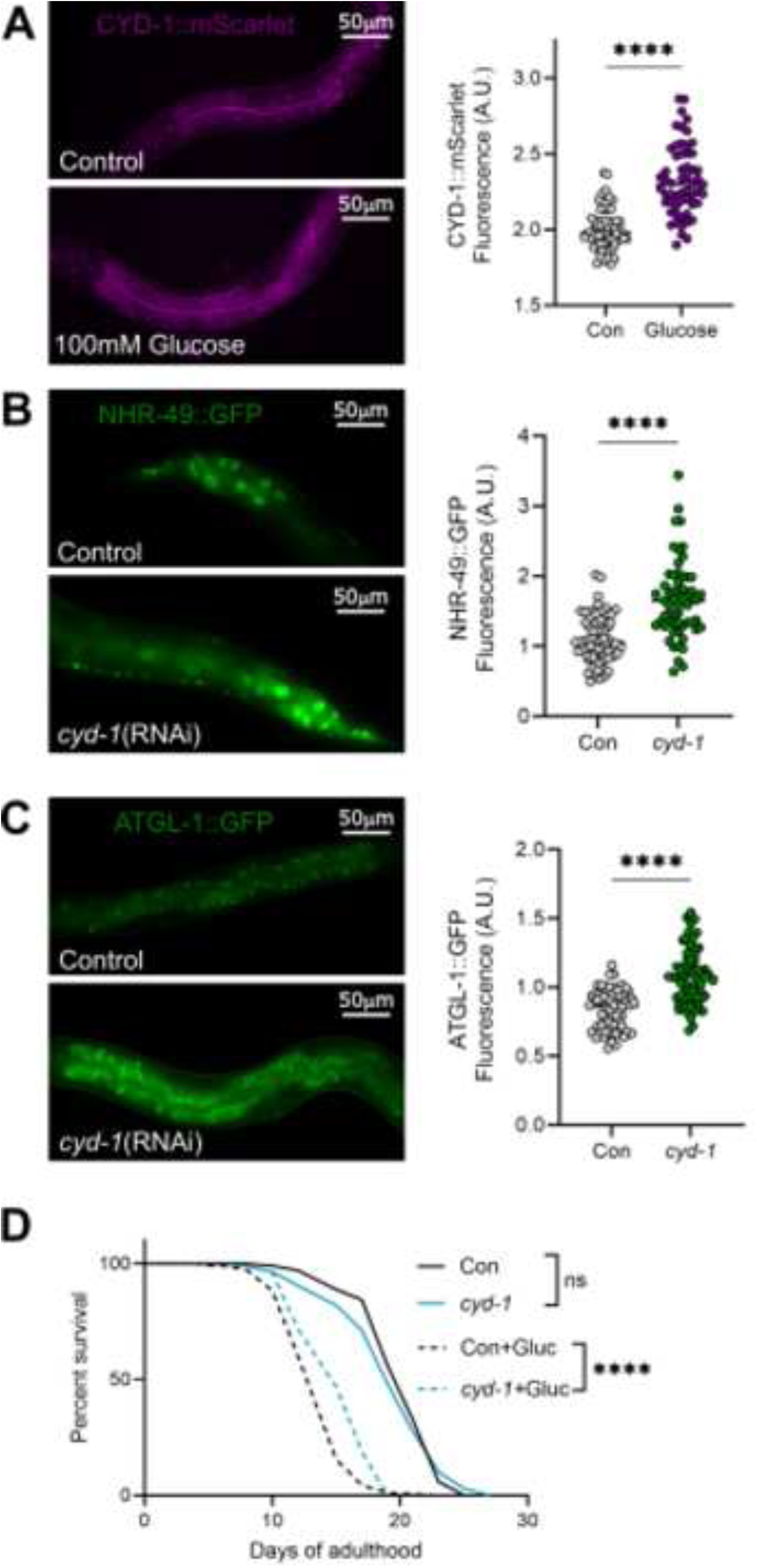
The role of cyclin D1 (Cyd-1) in the response to high glucose feeding in *C. elegans*. (**A**) Increased CYD-1::mScarlet expression in the intestine of L4 worms grown on 100mM glucose from eggs. (**B**) Increased NHR-49::GFP expression in the intestine (hindgut) of L4 animals in response to cyd-1 RNAi. (**C**) ATGL-1::GFP expression is increased in the intestine of L4 animals in response to cyd-1 RNAi. (**D**) Effect of cyd-1 RNAi on lifespan in the presence of 100mM glucose starting in adulthood. Log-rank test: ns – not significant, ****p≤0.0001. Identical results were obtained in 2 biological replicates. For A-C, representative images are 40x magnification with identical exposure time (scale bar shown) and quantitation data are pooled from 3 independent replicates. Welch’s t-test ****p≤0.0001.

Since CYD-1 inhibited the expression of two mediators that promote longevity in *C. elegans* (NHR-49 and ATGL-1 [73; 76; 77]), we examined whether knockdown of this protein regulates lifespan in the setting of overnutrition. Glucose treatment starting on day 1 of adulthood caused a substantial reduction in lifespan in worms (Fig. 5E), consistent with published studies [68–70]. Treatment with *cyd-1* RNAi starting on day 1 of adulthood did not affect lifespan in control worms, but it significantly increased lifespan in the setting of glucose treatment. Cumulatively, the data in Fig. 5 show that CYD-1 is induced by overnutrition and regulates key nutrient-mediated proteins in *C. elegans* intestine in a manner similar to cyclin D1 in hepatocytes, and that it shortens lifespan in this setting.

## 3. DISCUSSION

The main finding of this study is that cyclin D1 plays an unexpectedly broad role in hepatocytes in response to feeding, suggesting that it acts as a “hub” protein in hepatic physiology in the absence of substantial cell cycle progression. Although there is a wealth of literature on cyclin D1, relatively few studies have examined its roles outside of cell proliferation and cancer. The data presented here support the concept that cyclin D1 can act primarily as a metabolic regulator beyond its well-characterized role in the cell cycle. A particularly striking finding was that cyclin D1 markedly promotes the expression of APR proteins that can act as SASP and are involved in disease states. Hepatocyte cyclin D1 is induced in aged liver, premature aging, and MASLD – without commensurate proliferation – and thus may play a role in the pathophysiology of these conditions. Importantly, cyclin D1/CYD-1 regulates nutrient-mediated signaling and lifespan in *C. elegans*, indicating that these effects are highly conserved in evolution.

Our prior studies have shown that short-term cyclin D1 overexpression is sufficient to promote robust mitogen-independent hepatocyte proliferation and liver growth, even in the setting of TORC1 inhibition and protein deprivation [21–23; 78]. Compared to the quiescent state, substantial metabolic rewiring is required to support growth and proliferation through mechanisms that are incompletely characterized [79; 80]. In proliferating hepatocytes, cyclin D1 diminishes broad aspects of “homeostatic” metabolism (in part by repressing key transcription factors including HNF4α, PPARα, and ChREBP) [24–27], while at the same time enhancing biosynthetic pathways [28]. Thus, cyclin D1 is a pleiotropic metabolic regulator during cell cycle progression in hepatocytes.

Previous reports have shown that cyclin D1 is induced in the liver by feeding and inhibits two key aspects of glucose metabolism, gluconeogenesis [32; 33] and glycogen synthesis [24]. In the latter study, deletion of cyclin D1 induced a 3-fold increase in hepatic glycogen stores in D1^ΔHep^ mice refed a high-carbohydrate diet, indicating a substantial metabolic effect; similarly, in cultured AML12 hepatocytes, cyclin D1 siRNA promoted an equivalent increase in glycogen content, demonstrating that this is a cell-autonomous response. Interestingly, the induction of cyclin D1 in hepatocytes by a high-sucrose carbohydrate diet in this study has a parallel in the plant *Arabidopsis*, where sucrose induces D-type cyclin expression, suggesting that this is an evolutionarily conserved response [81]. The marked effect of cyclin D1 inhibition on glycogen synthesis led us to examine whether it more broadly regulates hepatic metabolism in response to feeding.

Based on RNA-seq analysis, cyclin D1 modulates a substantial portfolio of transcripts in response to feeding involved diverse hepatic functions. In refed liver, cyclin D1 knockout affects gene expression in a manner that significantly overlaps with the fasting response, such as reduced expression of APR/SASP and inflammatory mediators. On the other hand, some feeding-stimulated responses are enhanced in D1^ΔHep^ liver, including ChREBP-mediated lipogenic gene expression and glycogen synthesis [24]. Triglyceride levels in refed liver were lower in the absence of cyclin D1, consistent with our findings that this protein inhibits multiple steps of lipid catabolism [24; 25; 27]. Although further study is required to define mechanisms, these findings provide new insight into proliferation-independent roles of cyclin D1.

We were surprised to find that the most highly induced pathways by cyclin D1 in refed liver included many genes involved in APR/SASP and inflammatory signaling. In response to acute and chronic systemic stresses such as infection, inflammation, and dysmetabolic conditions, hepatocytes produce high levels of APR proteins that that play a role in microbial resistance, systemic immune function, metabolism, and tissue repair [6–10]. Prior studies have found that refeeding after a period of fasting induces expression of APR proteins and inflammatory markers in the liver in mice and humans [40–43], but the underlying mechanisms have not been well defined. Notably, numerous APR proteins have been shown to promote metabolic dysfunction and other deleterious effects in the liver and other organs, and are well-characterized SASP factors that promote aging-related pathologies [6–10].

For example, Lcn2 is a secreted multifunctional APR and SASP protein that is induced by feeding and remarkably responsive to cyclin D1 in our liver models; it is substantially diminished in refed D1^ΔHep^ liver and serum and was the most highly upregulated transcript following short-term cyclin D1 overexpression [34]. Although it is expressed in numerous cell types, hepatocytes are the main source of plasma Lcn2 in the setting of bacterial infection or liver injury [82]. This protein contributes to hepatic injury and fibrosis through activation of inflammatory and stellate cells [83; 84]. Increased circulating Lcn2 levels appear to promote metabolic dysfunction and related disorders including diabetes, insulin resistance, and neuroinflammation, including in the settings of MASH and aging [83–89]. Another example is PAI-1 (Serpine1), which is an extensively-studied APR/SASP protein. Although it is expressed in many cell types, hepatocytes are a major source of this protein [45; 46]. It is linked to obesity, the metabolic syndrome, MASLD, thrombotic disorders, and other diseases in humans, and studies in mice suggest that it plays a causative role in these conditions [44–48]. Furthermore, genetic deficiency of PAI-1 prevents premature aging in a mouse model [90], and is associated with longevity in humans [91].

We interpret the data from our studies to suggest the possibility that chronic expression of cyclin D1 might play a maladaptive role in hepatic metabolism and aging independently of its role in liver regeneration. Numerous aging-related pathologies are promoted by overnutrition via nutrient-regulated pathways, including MASH and other liver diseases [92; 93]. Conversely, the best characterized intervention to promote longevity is calorie restriction [4; 5]. We now show that cyclin D1 plays a significant role in feeding-stimulated liver physiology, and that knockout of this protein has overlapping effects with fasting. Hepatic expression of cyclin D1 is induced in aging; in the *Ercc1*-deficient mouse model of accelerated aging (which has marked liver dysfunction [61]), most hepatocytes are cyclin D1-positive despite the absence of proliferation. A recent study using immortalized human hepatocytes found that chronic hyperinsulinemia induced cyclin D1 along with senescence; furthermore, in bulk RNA-seq of human livers with MASLD, increased cyclin D1 expression correlated with higher expression of several senescence-associated genes [64]. Although a causative role has yet to be established, our current and prior studies indicate that cyclin D1 has actions in hepatocytes that are commonly observed in aging and senescence [4; 13; 15; 92–96], including: *(1)* Induction of SASP and inflammatory mediators, *(2)* increased p300-mediated gene expression, *(3)* decreased HNF4α and PPARα activity (and thus diminished homeostatic metabolic function) [24–26], *(4)* diminished lipolysis and fatty acid oxidation [24; 25; 27], *(5)* reduced autophagy [27], *(6)* increased aneuploidy and polyploidy [97], *(7)* increased p21 expression [22], and *(8)* enhanced cellular anabolic metabolism [28]. Future studies, possibly including the use of D1^ΔHep^ mice, are warranted to ascertain whether cyclin D1 plays a pathophysiologic role in liver aging, senescence, and metabolic diseases.

To further examine the potential biologic importance of cyclin D1 in feeding and aging, we took advantage of the *C. elegans* model, which has been extensively utilized to discover pathways involved in metabolism, lifespan, and other fundamental processes relevant to human diseases. Basic results from our fast-refeeding studies in the liver had analogous findings in *C. elegans* intestine, which carries out “liver” functions in this organism [66]: Cyclin D1/CYD-1 is induced with excess carbohydrate feeding, and it regulates the expression of key nutrient-mediated proteins. Furthermore, in young adult worms (where there is no longer cell proliferation), inhibition of CYD-1 prolonged lifespan in the setting of overnutrition. Future studies in *C. elegans* will provide insight into cell cycle-independent roles of cyclin D1 in nutrient-mediated signaling, metabolism, and aging.

In summary, cyclin D1 broadly regulates the hepatic response to feeding, which appears to be conserved in evolution. It is required for the induction of APR/SASP proteins in this setting, and thus may play an unexpected cell cycle-independent role in innate immunity, inflammation, hepatic protein and hepatokine production, and other key aspects of liver biology. Expression of this protein in hepatocytes is induced in aging and MASLD without commensurate proliferation, and thus it may regulate physiology in these settings. These studies provide further evidence that metabolic regulation is an important function of cyclin D1.

## ACKNOWLEDGEMENTS

We thank Wendy Larson for performing immunohistochemical studies, Peter Sicinski for providing cyclin D1^fl/fl^ mice, Shannon Jannatpour for help procuring liver biopsy specimens, and the Peter Adams lab for helpful discussion. This work utilized the computational resources of the NIH HPC Biowulf cluster (http://hpc.nih.gov).

## CRediT AUTHORSHIP CONTRIBUTION STATEMENT

H. Wu, J. Hauser, N. Yang, N. Timchenko, M. Klaers, R. Salekeen, J. Manivel, J. Abrahante, L. Laux, M. Yousefzadeh, M. Schonfeld, I. Tikhanovich, O. Adeyi: Methodology, Investigation, Formal analysis, Data curation. H. Wu, M. Yousefzadeh: Conceptualization. N. Timchenko, I. Tikhanovich, S. Ikramuddin, S. Monga, L. Niedernhofer, P. Sen, M. Gill, J. Albrecht: Conceptualization, Funding acquisition, Project administration. J. Albrecht: Writing - original draft, Writing - review & editing. M. Gill: Writing - review & editing.

## FUNDING

This work was supported by the following NIH grants:

DK54921 (J. H. A.), R01DK062277 (S.P.M.), CA278834 (N.T.), AA031270 (I.T.), R21DK122832 (S.I.), R01DK128325 (S.I.)

## DATA AVAILABILITY

Data will be made available on request. Sequencing data will be available in Gene Expression Omnibus (GEO).

## APPENDICES. SUPPLEMENTARY DATA

Supplementary Material (PDF)

Supplementary Tables S1 and S3 (spreadsheets)

