## Supplemental Material for "Cyclin D1 regulates the hepatic response to feeding: Evidence for non-cell cycle roles in the liver"

**Supplementary material**

**Table S1: RNA-seq data for refed D1^ΔHep^ and control liver** (enclosed in a separate file)

**Table S2: The top IPA canonical pathways in refed D1^ΔHep^ liver** (relative to control refed):

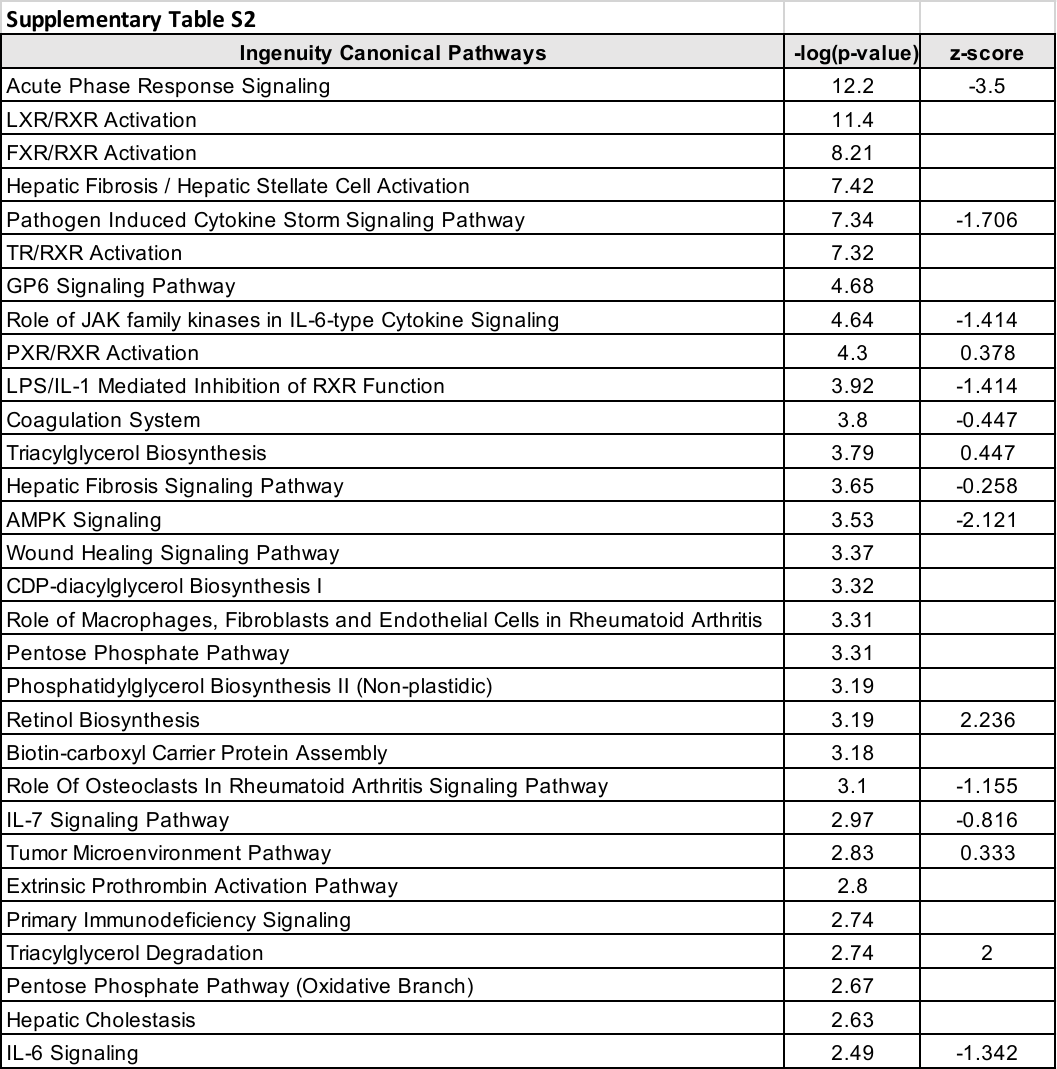

**Table S3: Transcripts in refed D1^ΔHep^ liver that were oppositely regulated by acute cyclin D1 overexpression** (enclosed in a separate file)

**Supplementary Figure S1**: **Regulation of ChREBP-mediated genes by cyclin D1 in refed liver**. (**A**) The most highly regulated transcription factors in refed D1^ΔHep^ livers (relative to refed control) predicted by IPA Upstream Regulator analysis (z-score of +/-2 or greater). (**B**) Fold-change in ChREBP-regulated genes in refed D1^ΔHep^ livers (relative to refed control). (**C**) Western blot of ChREBP and the lipogenic enzyme Acaca (ACC). (**D**) Phosphorylation of ChREBP by recombinant cyclin D1/Cdk4. An adenovirus expressing ChREBP-Flag was used to overexpress this protein in liver, which was harvested after an overnight fast. ChREBP-Flag was isolated and purified using anti-FLAG beads are previously described [1]. This was incubated with cyclin D1/Cdk4 and ATPγS as previously described [1], followed by western blot with anti-thiophosphate ester and ChREBP antibodies.

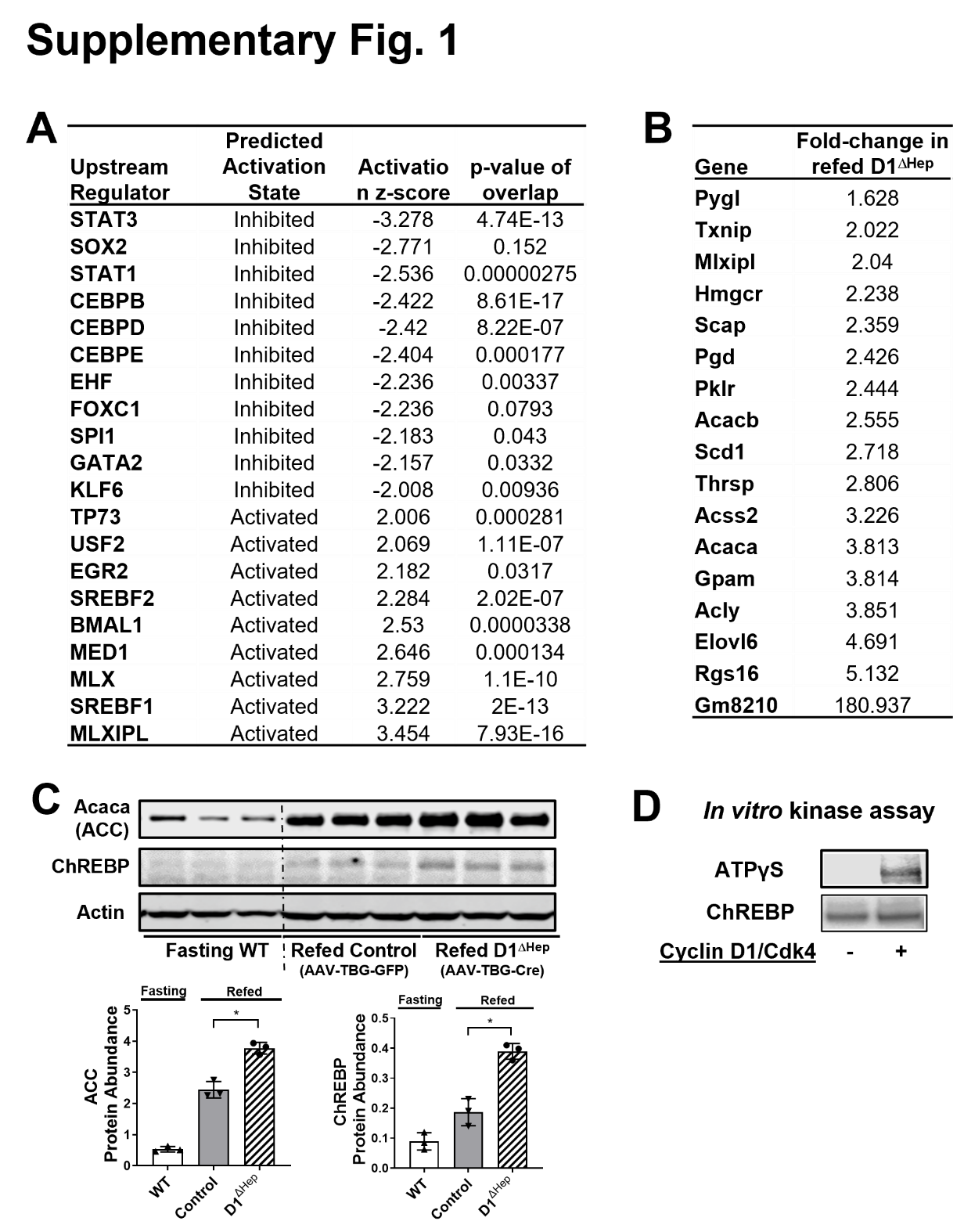

**Supplementary Figure S2**: **Mouse liver immunohistochemistry**. Some of the cyclin D1 IHC images are also shown in Fig. 4, but are included her for ease of viewing. (**A**) Regenerating young mouse liver 42 h after 2/3 partial hepatectomy (PH) (as in ref. [2]) showed induction of cyclin D1 as well as proliferation as shown by ph-Histone H3 in hepatocytes. Aged mouse liver showed increased hepatocyte cyclin D1 expression in Zone 2 but no proliferation. (**B**) In Ercc1^-/Δ^ liver at 6 months, most hepatocytes showed cyclin D1 expression but no proliferation was observed.

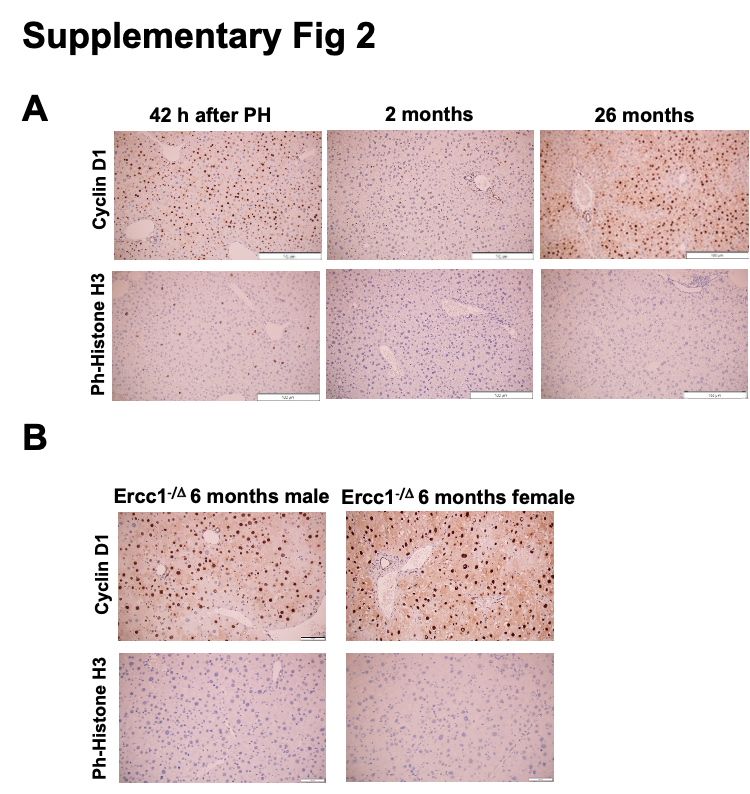

**Supplementary Figure S3: Human liver immunohistochemistry**. Some of the cyclin D1 IHC images are also shown in Fig. 4, but are included her for ease of viewing. (**A**) Histologically normal liver from young and aged human liver (3 individuals in each group). Aged liver had increased cyclin D1 expression without proliferation (as evidenced by ph-Histone H3 expression) (**B**) Hepatocyte cyclin D1 is upregulated in MASLD/MASH without associated cell proliferation. A specimen with autoimmune hepatitis showed numerous cyclin D1-positive hepatoctyes with associated proliferation as evidenced of Ki-67 and ph-Histone H3 immunostaining. Patients with MASLD/MASH had numerous cyclin D1-expressing hepatocytes without proliferation (some immune cells were positive for Ki-67 and ph-Histone H3).

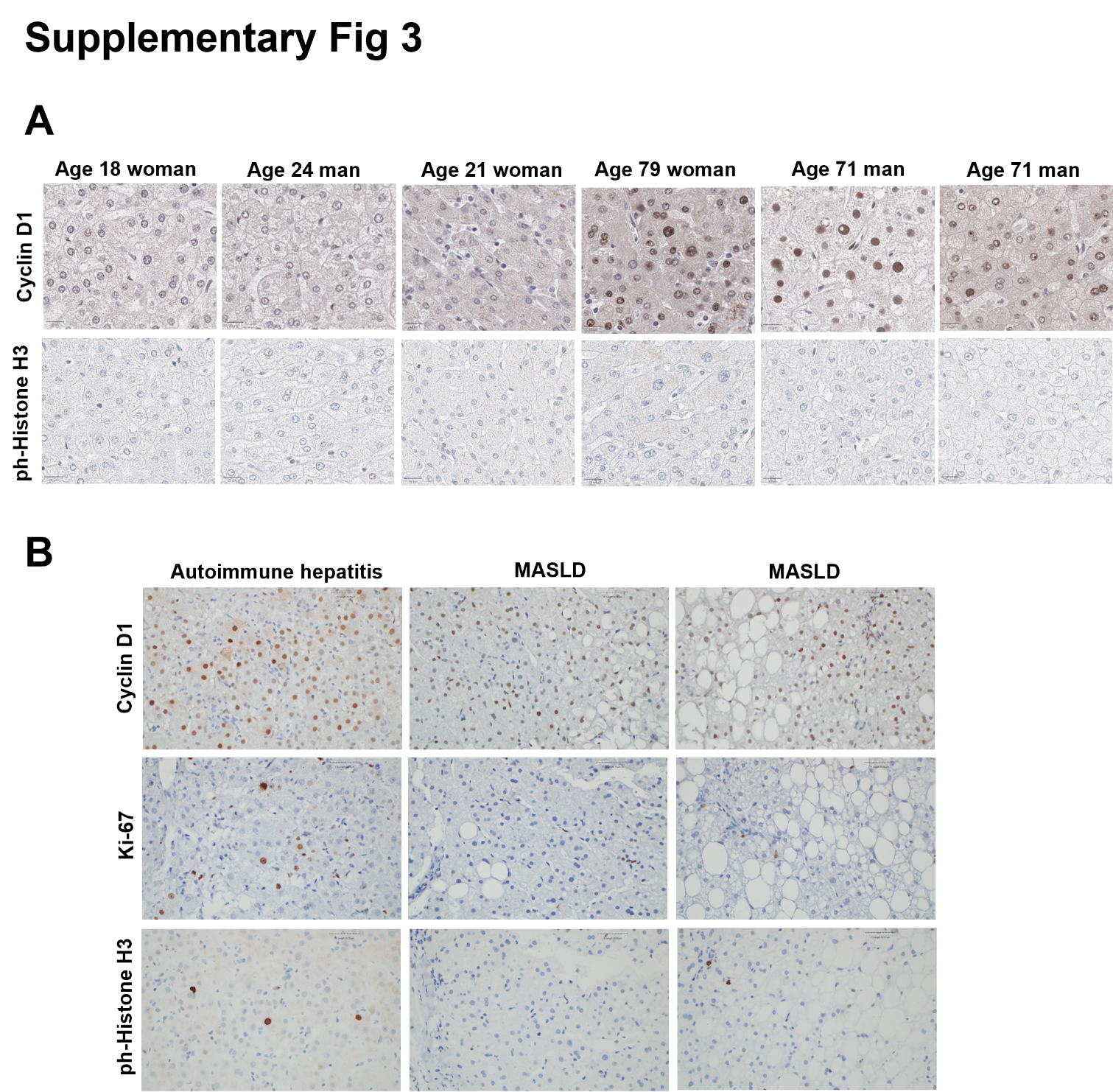

**MATERIALS AND METHODS**

**Mouse studies**

Mouse studies were approved by the institutional animal use and care committees at the Minneapolis VA Health Care System, the University of Minnesota, and the University of Kansas Medical Center. Hepatocyte-specific cyclin D1 knockout mice (D1^ΔHep^) were generated as previously described [24]. Briefly, one week before experiments, male Cyclin D1^fl/fl^ mice [36] were injected intravenously with 2 x 10^11^ viral particles of either the control vector adeno-associated virus serotype 8 (AAV8)-thyroxine-binding globulin (TBG)-enhanced green fluorescent protein (GFP) (AAV8-TBG-GFP, the control vector) or with AAV8-TBG-CRE (Addgene) to achieve acute hepatocyte-specific knockout as previously described [37; 38].  CEBPB^fl/fl^ mice were similarly treated to obtain hepatocyte-specific knockout (CEBPB^ΔHep^) mice [3]. At 8-10 weeks of age, mice were fasted for 20 h, or fasted and then refed a high-carbohydrate diet (88122i, Envigo) for 24 h prior to harvest as previously described [24; 26]. Tissue from young female liver taken 42 h after 2/3 partial hepatectomy was obtained as previously described [2]. Liver tissue was frozen in liquid N_2_ and stored at -80 or fixed in formalin for paraffin embedding and microscopy.

Tissue from young (2-6 month) and aged (26-27 month) FVB:C57Bl6 livers were used for IHC and western blot.  Ercc1-deficient (Ercc1^-/Δ^) mice and corresponding age-matched WT mice were bred as previously described [39]. For western blot, Ercc1^-/Δ^ mice were harvested at age 16 weeks, and WT mice were harvested at 18-19 weeks; for IHC, these livers were harvested at age 6 months.

**Human liver specimens**

Deidentified human liver biopsy specimens from obese patients with MASLD taken prior to and one year after successful weight loss after vertical sleeve gastrectomy were obtained from a prior study at the University of Minnesota (NCT03997422). Archival liver biopsy specimens determined to be histologically normal from young and aged adults were obtained from the University of Pittsburgh Liver Institute Clinical Biospecimen Repository and Processing Core.

**Western blot**

Liver tissue was homogenized and lysed in RIPA buffer with protease and phosphatase inhibitors as previously described [2]. Western blot was performed as described using 50 mg of protein was used per lane [2].

**Nuclear protein studies**

Liver was homogenized in buffer A containing 10mM tris-HCl, ph 7.5; 30mM NaCl; 5mM EDTA, 5mM b-mercapto-ethanol  and 10% glycerol. After centrifugation at 10,000 rpm (cold room) for 10 min, the pellet was incubated with buffer B (the same as buffer A, but containing 0.42M NaCl) for 20 min. After centrifugation at 10,000 rpm (10 min), the supernatant (nuclear extract, NE) was collected. For immunoprecipitation, the NE was diluted twice by 10mM Tris HCL followed by immunoprecipitation with antibody to cyclin D1 NE and immunoprecipitants were subjected to western blot as previously described [4].

**Antibodies**

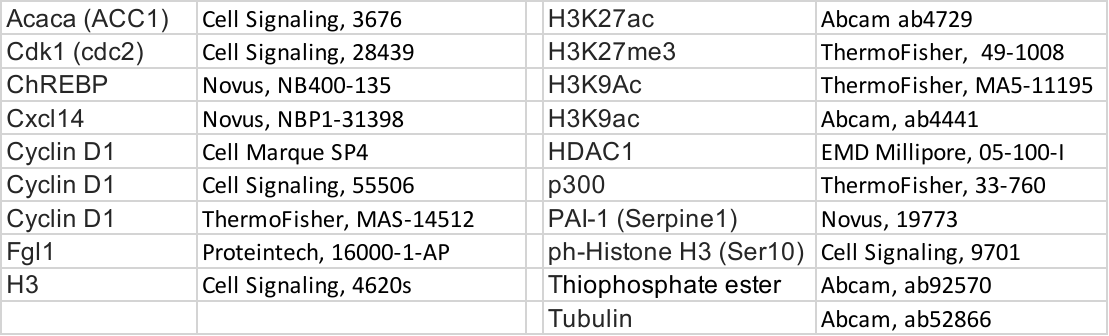

**ELISA**

Serum collected from mice was analyzed for ELISA using kits from R&D systems according to the manufacturer’s instructions (Mouse Lipocalin 2 DY1857; Mouse Serpin E1/PAI-1 DY3828-05).

**RNA-Sequencing (RNA-seq) and differential expression analysis**

Total RNA was isolated from liver and quantified using a fluorimetric RiboGreen assay (Thermo Fisher). Total RNA samples were converted to Illumina sequencing

libraries using Illumina’s TruSeq RNA Sample Preparation Kit. mRNA from a normalized input mass of total RNA was isolated using oligo-dT coated magnetic beads, fragmented and then reverse transcribed into cDNA. Libraries were amplified using 15 cycles of PCR. Final library size distribution was validated using capillary electrophoresis and quantified using fluorimetry (Pico Green) and via Q-PCR. The libraries were sequenced on the Illumina NovaSeq platform. Base call (.bcl) files for each cycle of sequencing were generated by Illumina Real Time Analysis (RTA) software. Quality control, data alignment, and gene quantification were analyzed using the CHURP pipeline [5] at the University of Minnesota Supercomputing Institute. 2 x 150bp FASTQ paired-end reads for 24 samples (26.3 million reads average per sample) were trimmed using Trimmomatic (v0.33) enabled with the optional “-q” option; 3bp sliding-window trimming from 3’ end requiring minimum Q30. Quality control on raw sequence data for each sample was performed with FastQC. Read mapping was performed via HISAT2 (Kim et al., 2019) (v2.1.0) using the mouse genome (GRm38) as reference. Gene quantification was performed using FeatureCounts for raw read counts. Differentially expressed genes were identified using the EdgeR (negative binomial, R programming) feature in CLCGWB (Qiagen) using raw read counts. We filtered the generated list for a minimum 1.5X absolute fold change and FDR-corrected <0.05. The transcripts regulated on this list were analyzed were analyzed through the use of IPA (Ingenuity Systems, Qiagen) as previously described [2].

**Chromatin immunoprecipitation (ChIP) and RT-PCR**

ChIP was performed as previously described [4; 6]. Primers used for PCR of the *Lcn2* genomic region were: Reverse 5’ tgggcctacctggctgccagcc -3, Forward 5’-gggaatgtccctctggtccccc- 3’. Polymerase chain reaction conditions were as follows: 99°C for 5 minutes, followed by 34 cycles of 94°C for 30 seconds, 60°C for 1 minute, 72°C for 30 seconds, and a final extension step at 72°C for 10 minutes. The polymerase chain reaction products were analyzed by 6%–8% polyacrylamide gel electrophoresis.

**ChIP-sequencing (ChIP-seq) of liver tissue**

H3K9ac, H3K27ac, and H3 ChIP were performed using the SimpleChIP plus sonication kit (Cell Signaling, 56383) following manufacturer’s instructions. Briefly, ~1000 mg of tissue was minced and crosslinked in 1% formaldehyde for 30 min at room temperature, followed by quenching with glycine and washing in ice-cold PBS. Tissue was lysed and homogenized, and nuclei were isolated and resuspended in nuclear lysis buffer. Chromatin was fragmented by sonication using a Covaris S220 to generate DNA fragments predominantly <500 bp. The lysate was clarified by centrifugation, and an aliquot was reserved as input. Equal amounts of chromatin (10 µg) were incubated with specific antibodies overnight at 4°C, followed by capture with Protein G magnetic beads. Beads were extensively washed under low- and high-salt conditions, and bound chromatin was eluted and reverse crosslinked at 65°C. DNA was then purified using spin columns and used for downstream analysis. DNA (~5 ng) was used to prepare libraries with the NEBNext Ultra II library preparation kit with unique dual index primers (New England Biolabs). The library quality and quantity were verified by BioAnalyzer DNA 1000 (Agilent) run and qPCR with NEBNext Library Quant kit (New England Biolabs) respectively. The liver libraries were pooled and paired-end sequenced on the NextSeq 2000 platform (Illumina) using the P3 100 cycle kit.

Sequencing reads (~25 million paired end reads per sample) were processed using a standardized pipeline. The reads were first de-multiplexed generating compressed FASTQ files by the on-board DRAGEN informatics pipeline (Illumina DRAGEN FASTQ Generation/v3.7.4) on the NextSeq 2000. FASTQ reads were concatenated across lanes and adapter trimming was performed with Trim Galore (trimgalore/v0.6.7), followed by quality control using FastQC (fastqc/v0.11.9) and summary reporting with MultiQC (multiqc/v1.9). Reads were aligned to the mm10 genome using Bowtie2 (bowtie/2) using the end-to-end parameter. Aligned reads were filtered and processed with SAMtools (samtools/v1.17) and Sambamba (sambamba/v1.0.0). PCR duplicates were removed using Picard (picard/v3.1.0). ENCODE blacklisted genomic regions were excluded using BEDTools (bedtools/v2.30.0). Biological replicates were merged at the BAM level using SAMtools (samtools/v1.17) to generate condition-specific datasets for each histone modification and H3 control. Merged BAM files were indexed and converted to normalized genome coverage tracks (bigWig) using deepTools (deeptools/v3.5.0) with RPKM normalization. To control for background signal, differential tracks were generated using deepTools (deeptools/v3.5.4) by subtracting H3 from histone modification tracks (bigwigCompare), producing normalized enrichment profiles for visualization as previously described [7].

**Immunohistochemistry (IHC)**

Sections of formalin-fixed paraffin-embedded tissue sections were stained with cyclin D1, ph-Histone H3, and Ki-67 as previously described [2; 8].

**C. elegans studies**

*C. elegans* strains were maintained as previously described [9]**.** Bristol N2 (wild-type), JJ1271[*glo-1(zu391)*], PMD150[*utsls4(nhr-49p::nhr-49::GFP + myo-2p::mCherry)*] and VS20[*hjIs67(atgl-1p::atgl-1::GFP + mec-7::RFP)]* were obtained from the *Caenorhabditis* Genetics Center (University of Minnesota).

To generate an endogenous *cyd-1* translational reporter, we used CRISPR / Cas9 genome editing to introduce mScarlet::AID::FLAG into the *cyd-1* genomic locus [10]. A crRNA targeting a protospacer adjacent motif (PAM) sequence close to the 3’ end of the *cyd-1* coding sequence was identified using Wormbase [11] and a double-stranded DNA repair template was amplified from plasmid pJW2098 (a gift from Jordan Ward; Addgene plasmid # 163094 ; http://n2t.net/addgene:163094 ; RRID:Addgene_163094) [12]. Cas9, crRNA, and trans-activating crRNA (tracrRNA) were obtained from Integrated DNA Technologies (IDT). The injection mix was prepared according to Wang et al. [13]. Approximately 30 P0 animals were injected and singled onto individual plates. F1 roller animals were isolated from a single jackpot plate and F2 *cyd-1* animals were identified by the genotyping. One line with the correct edit was isolated and outcrossed three times to yield MGL554[*cyd-1(jlu37[cyd-1::mScarlet::AID::FLAG*])*.* MGL554 was then crossed into JJ1271 to generate MGL581[*cyd-1(jlu37); glo-1(zu391)*].

A *cyd-1* RNAi clone was obtained from the Ahringer RNAi library and sequenced to confirm the correct identity. RNAi bacteria were grown in the presence of tetracycline (10 µg/mL) and carbenicillin (50 µg/mL) antibiotics, and single colonies were inoculated in LB with carbenicillin (50 µg/mL) for 16–20 hours at 37°C. Nematode growth medium (NGM) plates containing 1 mM IPTG and 50 µg/ mL carbenicillin were seeded with an overnight culture of bacteria, dried in a sterile hood, and maintained at room temperature for 48 h. Ten to fifteen gravid adults were transferred to fresh RNAi plates, allowed to lay eggs for 2 hours, and removed. The L4440 empty vector was used as a negative control.

Eggs from a synchronous lay were grown on RNAi bacteria for 48h and imaged at the L4 stage. Images were obtained using a Hamamatsu ORCA-Fusion BT camera on a Nikon Eclipse Ti2-E inverted microscope with Nikon CFI Plan APO 𝛌D 40X or 60x(oil) objectives, illuminated with a multichannel Nikon D-LEDI fluorescence LED illumination system which maintains constant intensity and light source alignment. Images were compiled and analyzed with Nikon NIS Elements software. All representative images were generated using a pseudocolor merge function, consistent thresholding based upon the background, and deconvolution projection through the NIS Elements.

Lifespan assays were performed at 20°C on RNAi plates with or without 100mM glucose and 12.5μg/mL 5-fluoro-2′-deoxyuridine (FUDR) starting at the first day of adulthood. Animals were transferred again on the second day of adulthood, then every 2-3 days until completion. Death was scored by loss of touch-provoked movement, and animals lost due to uterine prolapse or crawling up the side of the petri dish were censored.

**Statistical analysis**

Replicates shown in the figures represent independent experiments done in parallel. Data are expressed as mean ± SD. Statistical analysis was performed using GraphPad software (GraphPad Software, Inc; www.graphpad.com). Comparisons between two groups were made by Student t test, and significant differences were noted (*p < 0.05; **p < 0.01; and ***p <0.001).
